# Mitochondrial H2S and cytochrome c coordinate guard cell redox dynamics during stomatal immunity

**DOI:** 10.64898/2026.11.20.768174

**Authors:** Rosario Pantaleno, Denise Scuffi, Florencia P. Coronel, Elina Welchen, Carlos García-Mata

## Abstract

Stomatal pores modulate gas exchange between plants and the environment and are the first layer of defense against pathogens entry. Hydrogen sulfide (H_2_S) is a recognized component of the signalling network in guard cells that control stomatal responses. We previously showed that the mitochondrial H_2_S source β-cyanoalanine synthase (CAS-C1) is required for flg22-induced stomatal closure. Using genetically encoded redox biosensors in *Arabidopsis thaliana* guard cells, we found that the mitochondrially-targeted H_2_S donor AP39 and the pathogen-associated molecular pattern (PAMP) flg22 both increased mitochondrial H_2_O_2_ within 5 min, while the flg22-triggered rise was attenuated in *cas-c1* mutants. Dissipating the mitochondrial membrane potential (Δψm) with the uncoupler 2,4-dinitrophenol prevented both mitochondrial oxidation and stomatal closure. Cytochrome c (CYTc), the mobile electron carrier between complexes III and IV, proved essential for this pathway. CYTc-deficient lines showed a stomatal response similar than wild-type (wt) to ABA and light but failed to trigger stomatal closure upon exposure to different PAMPs. In these plants, neither AP39 nor flg22 induced guard cell mitochondrial hyperpolarization or an increase in cytosolic H_2_O_2_. By the dimedone-switch method, we show that CYTc is persulfidated *in vivo*, identifying this respiratory protein as a target of H_2_S-mediated posttranslational modification. At the whole-plant level, *cytc* plants were more susceptible to spray inoculation with *Pseudomonas syringae pv. tomato hrcC*⁻ at early stages of infection. Together, these results position CYTc as a mitochondrial node that links H_2_S signalling to guard cell redox dynamics during stomatal immunity.

## Introduction

Plant pathogens severely affect crop yield and quality, generating substantial economic losses and threatening global food security goals. Anthropogenic activities and global climate change shape phytopathogen distributions by driving fluctuations in atmospheric CO_2_ concentrations, temperature, and water availability. To face biotic challenges, plants have developed sophisticated defense mechanisms, including rapid stomatal closure as an early physical barrier that limits pathogen entry and subsequent infection (Melotto et al., 2024).

Stomata serve as critical gates that regulate the trade-off between carbon fixation and water loss through the transpirational stream (Lawson and Matthews, 2020). These microscopic pores are delimited by pairs of specialized guard cells in the epidermis of the aerial tissues of most land plants. Detection of pathogen-associated molecular patterns (PAMPs) by specific plasma membrane pattern-recognition receptors (PRRs) initiates a subset of early defense responses known as pattern-triggered immunity (PTI) (DeFalco and Zipfel, 2021), which includes the activation of different signalling pathways in guard cells that induce stomatal closure within the first hour after exposure (Melotto et al., 2006). This response, called stomatal immunity, restricts pathogen entry to promote plant survival (Hou et al., 2024; Melotto et al., 2024). However, pathogens that overcome this barrier deliver virulent molecules within host cells (Yuan et al., 2021). Intracellular receptors recognize these effectors, eliciting effector-triggered immunity (ETI). Independent studies have shown that pathogen effectors induce stomatal closure to promote apoplastic water soaking, thereby favoring plant colonization, highlighting the fundamental role of stomatal movement regulation across layers of plant immunity (Hu et al., 2022; Roussin-Léveillée et al., 2022).

Guard cells detect, transduce, and integrate internal and external cues into a network of interconnected signalling pathways that control stomatal movement. Activation of these pathways ultimately modifies guard-cell turgor and volume to adjust stomatal pore size (Kim et al., 2010; Jezek and Blatt, 2017). In recent years, much progress has been made in describing both the structure and function of this network; however, new signalling players continue to emerge. Among them, the gasotransmitter hydrogen sulfide (H_2_S) has been pinpointed as a key regulator of stomatal closure, through an intricate crosstalk with several components from the signalling network, including ion channels, nitric oxide (NO), phosphatidic acid (PA), reactive oxygen species (ROS) and others (García- Mata and Lamattina, 2010; Scuffi et al., 2014; Papanatsiou et al., 2015; Jin et al., 2017; Scuffi et al., 2018; Chen et al., 2020; Shen et al., 2020; Pantaleno et al., 2021; Zhou et al., 2021).

At physiological pH, H_2_S coexists with its deprotonated form HS^−^; it acts as a reductant and nucleophile that reacts with oxidized cysteine residues (sulfenic acids) to form persulfides, a process known as protein persulfidation. Persulfidation is regarded as the main mechanism through which H_2_S exerts its biological function (Filipovic et al., 2018; Benchoam et al., 2019). This posttranslational modification modulates the enzymatic activity and localization of several target proteins. In plants, H_2_S has been reported to regulate physiological processes like autophagy, response to hypoxia, and stomatal closure (Scuffi et al., 2014; Chen et al., 2020; Laureano-Marín et al., 2020; Shen et al., 2020; Aroca et al., 2021; Zhou et al., 2021; Telara et al., 2025). In *Arabidopsis thaliana,* H_2_S can be enzymatically produced from cysteine or sulfite in different subcellular compartments (Aroca et al., 2020). The mitochondrial enzyme β-cyanoalanine synthase (CAS-C1) catalyzes the reaction between hydrogen cyanide and cysteine to produce ß-cyanoalanine and H_2_S, being the only characterized mitochondrial H_2_S (mtH_2_S) source (Yamaguchi et al., 2000; García et al., 2010). We recently reported that CAS-C1 participates in the regulation of stomatal immunity. Arabidopsis *cas-c1* mutants are unable to induce stomatal closure upon treatment with the PAMP flg22, a synthetic peptide from bacterial flagellum, while they are also more susceptible to spray-based infection with the bacterial strain *Pseudomonas syringae* pv. *tomato* DC3000 (Pantaleno et al., 2025). These findings position mitochondrial CAS-C1 as a novel regulatory component of the stomatal immune response; however, the role of mitochondrial-derived signals in the network that controls stomatal immunity remains largely overlooked.

PAMP recognition triggers a rapid increase in ROS levels in guard cells (Qi et al., 2018). Genetically encoded fluorescent sensors enable the real-time measurement of intracellular ROS dynamics within specific subcellular compartments, providing novel insight into their *in vivo* behavior (Wagner and Meyer, 2025). Using these biosensors, we reported that general and mitochondrially-targeted H_2_S donors produce a consistent hydrogen peroxide (H_2_O_2_) increase and the oxidation of glutathione redox potential (*E*_GSH_) in the cytosol of guard cells, indicating an interplay between H_2_S and the regulation of cellular redox state (Scuffi et al., 2018; Pantaleno et al., 2023).

Mitochondria not only produce energy to sustain most cellular demands, but they are also a major source of ROS (mitochondrial ROS; mtROS) (Møller, 2001). Electrons are transported through the mitochondrial Electron Transport Chain (mETC) to reduce molecular oxygen, establishing an electrochemical gradient in the inner mitochondrial membrane (Δψm) that drives ATP synthesis (Braun, 2020). While normal respiration generates superoxide, which is rapidly converted into H_2_O_2_, this mtROS production can be markedly increased upon certain stresses, including pathogen exposure (Bestwick et al., 1997; Cvetkovska and Vanlerberghe, 2012; Huang et al., 2016).

Plant mitochondria participate actively in the regulation of defense responses (Colombatti et al., 2014; Wang et al., 2022). Fungal and oomycete infection increases mtROS to initiate immune responses (Fuchs et al., 2016; Yang et al., 2025). Similarly, Arabidopsis leaf discs treated with flg22 exhibited a sustained rise in mitochondrial H_2_O_2_ levels, indicating that PAMP perception also affects mitochondrial redox state (Nietzel et al., 2019; Arnaud et al., 2023a). We recently showed that flg22-driven increase of cytosolic H_2_O_2_ is diminished in *cas-c1* mutants, pointing to a link between mtH_2_S and redox regulation during stomatal immunity through an underlying mechanism that remains to be elucidated (Pantaleno et al., 2025).

In animals, mtH_2_S has a cytoprotective effect, promoting mitochondrial bioenergetic activity through two main mechanisms. On one hand, H_2_S stimulates oxidative phosphorylation and ATP production by persulfidation of ATP synthase (Módis et al., 2016). On the other hand, H_2_S serves as an inorganic substrate for the respiratory chain, either by its oxidation followed by the delivery of electrons to ubiquinone, which are then transferred to mETC at complex III level, or by the direct reduction of complex IV or cytochrome c (CYTc) (Szabo et al., 2014; Murphy et al., 2019). Conversely, at higher concentrations, sulfide inhibits complex IV, resulting in a biphasic bioenergetic effect (Szabo et al., 2014). While experimental evidence on the potential effects of mtH_2_S on plant mitochondrial activity remains scarce, we recently reported that the mtH_2_S donor AP39 triggers dose-dependent stomatal closure and increases Δψm (Pantaleno et al., 2023). We also showed that mitochondrial complex IV activity and CYTc are required for AP39-dependent stomatal closure (Pantaleno et al., 2023).

CYTc is a small protein that delivers electrons to complex IV in the mETC and plays a key role in mitochondrial energy production (Welchen and Gonzalez, 2016). Interestingly, mammalian ferric CYTc oxidizes H_2_S, promoting protein persulfidation and enhancing the H_2_S-dependent signalling (Vitvitsky et al., 2018). While dedicated studies have analyzed the participation of CYTc in the regulation of diverse cellular processes, its involvement on stomatal immunity or its interplay with H_2_S has not been addressed so far (Racca et al., 2018; Florez-Sarasa et al., 2021; Racca et al., 2022; Canal et al., 2024; Coronel et al., 2025).

In this study, we present evidence that mtH_2_S and CYTc functionally regulate guard cell redox status during the flg22-induced stomatal immunity. We show that CYTc is required for PAMP-induced stomatal closure and for the mitochondrial and cytosolic redox responses that are triggered by flg22 and the mtH_2_S donor AP39 in guard cells. Endogenous mtH_2_S produced by CAS-C1 contributes to flg22-induced mitochondrial H_2_O_2_ accumulation, and CYTc is persulfidated *in vivo*. In line with this, CYTc-deficient plants are more susceptible to Pst hrcC⁻ at early stages of infection. We propose that CYTc links mitochondrial H_2_S signalling to guard cell redox changes during stomatal immunity.

## Materials and methods

### Plant Material and Chemicals

Seeds from *Arabidopsis thaliana* (Columbia-0 ecotype) wild type (wt), *cas-c1* mutants (SALK_022479 and SALK_068045C) and *cytc* double mutants (*1*b*2*a and *1*b*2*b) were available in our laboratory and previously described (Welchen et al., 2012; Racca et al., 2018; Pantaleno et al., 2025). Wt seeds expressing the biosensor mt-roGFP2-Orp1 were kindly provided by Prof. Schwarzländer (Nietzel et al., 2019), while *cas-c1* lines expressing this mitochondrial sensor were obtained by crossing. The F2 generation plants used were selected by fluorescence to verify the mt-roGFP2-Orp1 expression and by PCR-based genotyping to ensure homozygosity loci. *1*b*2*a lines expressing roGFP2-Orp1 were obtained by floral-dip transformation of mutated plants with *Agrobacterium tumefaciens* strain GV3101 carrying the roGFP2-Orp1 insert in a pH2GW7 vector under the control of a *CaMV35S* promoter. The vector was generated in the laboratory of Prof. Schwarzländer and kindly provided by the authors (Nietzel et al., 2019).

Plants were grown in a mixture of soil, perlite and vermiculite at a 3:1:1 ratio (v/v/v) at 20°C under short-day conditions (8-h-light/16-h-dark photoperiod, 200 µmol photons m^−2^ s^−1^), after stratification for 48 h at 4°C in the dark.

*Pseudomonas syringae* pv. *tomato* DC3000 wt and *hrcC^-^*strains were kindly provided by Dr. Georgina Fabro from CIQUIBIC-CONICET-UNC, Argentina. Bacteria were grown in King’s B agar medium (2% (w/v) proteose peptone, 1% glycerol, 0.15% (w/v) K2HPO4, pH 7.2) with 50 μg/mL rifampicin and 50 μg/mL kanamycin or rifampicin for 24 h at 28°C before infection.

Mitochondrial H_2_S donor AP39 (CAS N°: 1429173-57-8) was purchased in Cayman Chemical. Abscisic acid (ABA; CAS N°: 21293-29-8, horse-radish peroxidase (CAS N°: 9003-99-0), luminol (CAS N°: 521-31-3), 2,4-dinitrophenol (DNP, CAS N°:51-28-5), Protease Inhibitor Cocktail (N° P9599) and 4-Chloro-7-nitrobenzofurazan (NBF-Cl; N° 163260) were purchased in Sigma-Aldrich. High Capacity NeutrAvidin Agarose Resin (N° 29202) was purchased from Thermo Fisher Scientific™. DCP-Bio1 (CAS N° 882853-94-3) was purchased in Xoder Technologies, LLC. Tetramethylrhodamine methyl ester (TMRM) was purchased from Biotium. Flg22 (N-QRLSTGSRINSAKDDAAGLQIA-C) and elf18 (N-SKEKFERTKPHVNVGTIG-C) were purchased from ProteoGenix. Chitin oligosaccharide (NA-COS-Y) was bought from Yaizu Suisankagaku Industry Co., Ltd.

### Stomatal Aperture Measurements

Epidermal peels (epidermal strips) were excised from the abaxial side of 4–5-week-old A. *thaliana* wt or *cytc* leaves and immediately floated in opening buffer (OB: 5 mM MES-KOH, 50 mM KCl, pH = 6.1) for 1, 2, 3 or 4 h in the light to monitor light-induced stomatal opening, or 3 h followed by different treatments, as indicated in the figure legends, to assess stomatal closure (Pantaleno et al., 2024). To prepare *Pseudomonas* suspension, bacteria were grown at 28°C for 24 h in King’s B-agar plate with 50 μg/mL rifampicin and 50 μg/mL kanamycin or rifampicin and resuspended in OB until OD = 0.1 (λ = 600 nm). After treatments, stomata were digitized using an AmScope MU1000 camera coupled to an Olympus CKX53 microscope with a 40× lens (LUCPlanFLN, 0.6 numerical aperture). The stomatal aperture was measured using FIJI (ImageJ) analysis software (NIH, Bethesda, MD, USA). The stomatal aperture was quantified as the maximal width of the stomatal pore.

### Apoplastic ROS Burst Quantification

Leaf discs from 5-week-old Arabidopsis wt or *cytc* plants were floated in deionized water overnight. The following day, they were placed on a 96-well white plate containing a solution with 20 mM luminol and 0.02 mg mL−1 horseradish peroxidase. Leaf discs were treated or not with 100 nM flg22, and luminescence was measured immediately after the treatment in a luminometer (Thermo Scientific Luminoskan Ascent Microplate). The area under the curve (AUC) was calculated from measured relative light units (RLU) from each leaf disc.

### Confocal Laser Scanning Microscopy

Fluorescence images were acquired with a Nikon Eclipse C1 Plus confocal microscope.

Mitochondrial inner membrane polarization (TMRM): Mitochondria in GCs were imaged with a 100× lens (Nikon CFI Plan Apochromat Lambda, 1.45 numerical aperture, oil immersion). Epidermal peels from Arabidopsis thaliana wt or *cytc* leaves were floated in OB supplemented with 500 nM Tetramethylrhodamine methyl ester (TMRM) for 3 h in the dark and then treated with 100 nM AP39, 1 µM flg22 or 50 µM DNP as indicated in figure legends. TMRM was excited at 561 nm and emission was collected at 605/75 nm.

Redox-based sensors: Epidermal strips from wt, *cas-c1* or *cytc* Arabidopsis plants expressing either roGFP2-Orp1, mt-roGFP2-Orp1 or mt-Grx1-roGFP2, were floated in OB for at least 7 h under light for recovery as previously described (Scuffi et al. 2018) and then incubated with 100 nM AP39, 1 µM flg22, 50 µM DNP, or preincubated with DNP followed by AP39 or flg22 treatment, as indicated in each figure legend. To determine the biosensor’s spectral response range, we treated strips in each individual experiment with 10 min with 1 mM H_2_O_2_ and 20 mM dithiothreitol (DTT) for full oxidation and reduction, respectively. The epidermal peels were mounted, and images were collected with a 40×lens (Nikon CFI Super Fluor, 1.3 numerical aperture, oil immersion). Sensors were excited sequentially (Lambda frame) at 405 and 488 nm and emission was detected at 515/30 nm.

### Image Analysis

Images were analyzed employing FIJI (ImageJ) software (NIH, Bethesda, MD, USA). For fluorescence measurement from redox-based sensors, the background was subtracted from each channel and then the region of interest was delimited to either the cytosol (roGFP2-Orp1) or to the whole guard cell area (mt-roGFP2-Orp1 and mt-Grx1-roGFP2). We calculated the 405/488 nm ratio for each cell (Pantaleno et al., 2024). TMRM intensity was quantified by defining the region of interest in each guard cell area, followed by background subtraction. For all assays, fluorescence was measured as mean pixel intensity, and values were normalised to the wt control. The ratiometric false-colored images from guard cells expressing the redox-based sensors were obtained with FIJI software. Representative images were selected; the background was subtracted, and the images were thresholded. Finally, the images from each channel were divided using the Image Calculator, and a pseudocolor (Look Up Table Physics) was applied to the resulting image.

### Detection of protein persulfidation

*Arabidopsis thaliana* Col-0 seedlings were grown in liquid half-strength Murashige and Skoog (½ MS) medium supplemented with 1% (w/v) sucrose under long-day conditions (16-h-light/8-h-dark photoperiod, 200 µmol photons m^−2^ s^−1^). After 14 days, total proteins were extracted using lysis buffer (1× PBS pH 7.4, 1 mM EDTA, 2% (w/v) SDS, 1× Protease Inhibitor Cocktail. Persulfidated proteins were purified following the Dimedone-switch method (Aroca et al., 2022). Briefly, free thiols were blocked by incubation with 4-chloro-7-nitrobenzofurazan (NBF-Cl), and persulfides were selectively labeled with biotin-conjugated dimedone (DCP-bio1). Biotinylated proteins were then affinity-purified using NeutrAvidin agarose beads, and the eluted fraction was analyzed by Western blot.

### Western blot analysis

Sample preparation was performed using 15 µL of sample (I: Input, E: Eluted, and UF: unbound fraction) and 10 µL of 2× Laemmli buffer (375 mM Tris-HCl, pH 6.8, 9% SDS, 30% glycerol, 0.03 g bromophenol blue, and 9% β-mercaptoethanol), supplemented with a protease inhibitor cocktail (cOmplete; Roche) and 0.2 mM phenylmethylsulfonyl fluoride (PMSF). Proteins were denatured at 95°C for 5 min, then centrifuged at 12,000 × g for 2 min. Then, 20 µL of each sample was separated by 16% SDS-PAGE and transferred onto a polyvinylidene difluoride (PVDF) membrane. The membrane was incubated with a rabbit anti-CYTc antibody (AS08 343A, Agrisera; 1:2000 dilution). Detection was performed using horseradish peroxidase (HRP)-conjugated anti-rabbit IgG secondary antibodies (Agrisera; 1:50,000 dilution) and the SuperSignal™ West Pico PLUS chemiluminescent substrate (Thermo Fisher Scientific). After immunodetection, we stained the membranes with Coomassie Brilliant Blue R-250 (CBB) to confirm protein loading in each lane.

### Bacterial Spray-Inoculation

Five to 6-week-old Arabidopsis wt or *cytc* plants were sprayed with a *P. syringae pv. tomato* wt or *hrcC^-^*suspension (OD = 0.2 [λ = 600 nm] in 10 mM MgCl2, 0.02% (v/v) Silwet). Following spray inoculations, plants were kept covered with a transparent lid at 20°C under short-day conditions (8-h-light/16-h-dark photoperiod, 200 µmol photons m^−2^ s^−1^). Leaf discs samples were taken at Days 1 or 3 postinoculation (1 or 3 days postinoculation [dpi], respectively). Bacteria were growth in square Petri dishes with King’s B agar medium (2% (w/v) proteose peptone, 1% glycerol, 0.15% (w/v) K_2_HPO_4_, pH 7.2) with 50 μg/mL rifampicin and 50 μg/mL kanamycin or rifampicin for 48 h at 28°C. The number of colony-forming units (CFU) was determined after serial dilution and plating as described (Johansson et al. 2014).

## Statistical Analysis

Data analyses were performed using RStudio (R Foundation for Statistical Computing, Vienna, Austria; URL: https://www.R-project.org/). Statistically significant differences were analyzed using Generalized Linear Mixed Models (GLMM) with the lme function from the nlme package (Nonlinear Mixed Effects Models). Each independent experiment was included as a random factor. When multiple plant lines were compared, the genotype × treatment interaction was tested and included in the model. Tukey’s method was used for pairwise comparisons among genotypes, and Dunnett’s test was used to compare each treatment against the Control within each genotype, as indicated in the figure legends. The error distribution was assumed Gaussian and all effects were considered significant at p < 0.05.

## Results

### Mitochondrial H_2_S modulates mitochondrial redox state during stomatal immunity

To evaluate the crosstalk between mitochondrial redox dynamics and mtH_2_S, we used the biosensor mt-roGFP2-Orp1 to assess H_2_O_2_ changes in the mitochondrial matrix of guard cells (Nietzel et al., 2019). Epidermal peels were isolated from *Arabidopsis thaliana* wild-type (wt) plants expressing mt-roGFP2-Orp1 and treated with the mitochondrial-targeted H_2_S donor AP39 for different time periods. The fluorescence ratio 405/488nm was measured specifically in guard cells after incubation with the mtH_2_S donor. Results in Figure 1 show a consistent increase in mitochondrial H_2_O_2_ levels, which peaked 5 minutes after AP39 treatment (Fig. 1A).

**Figure 1.**
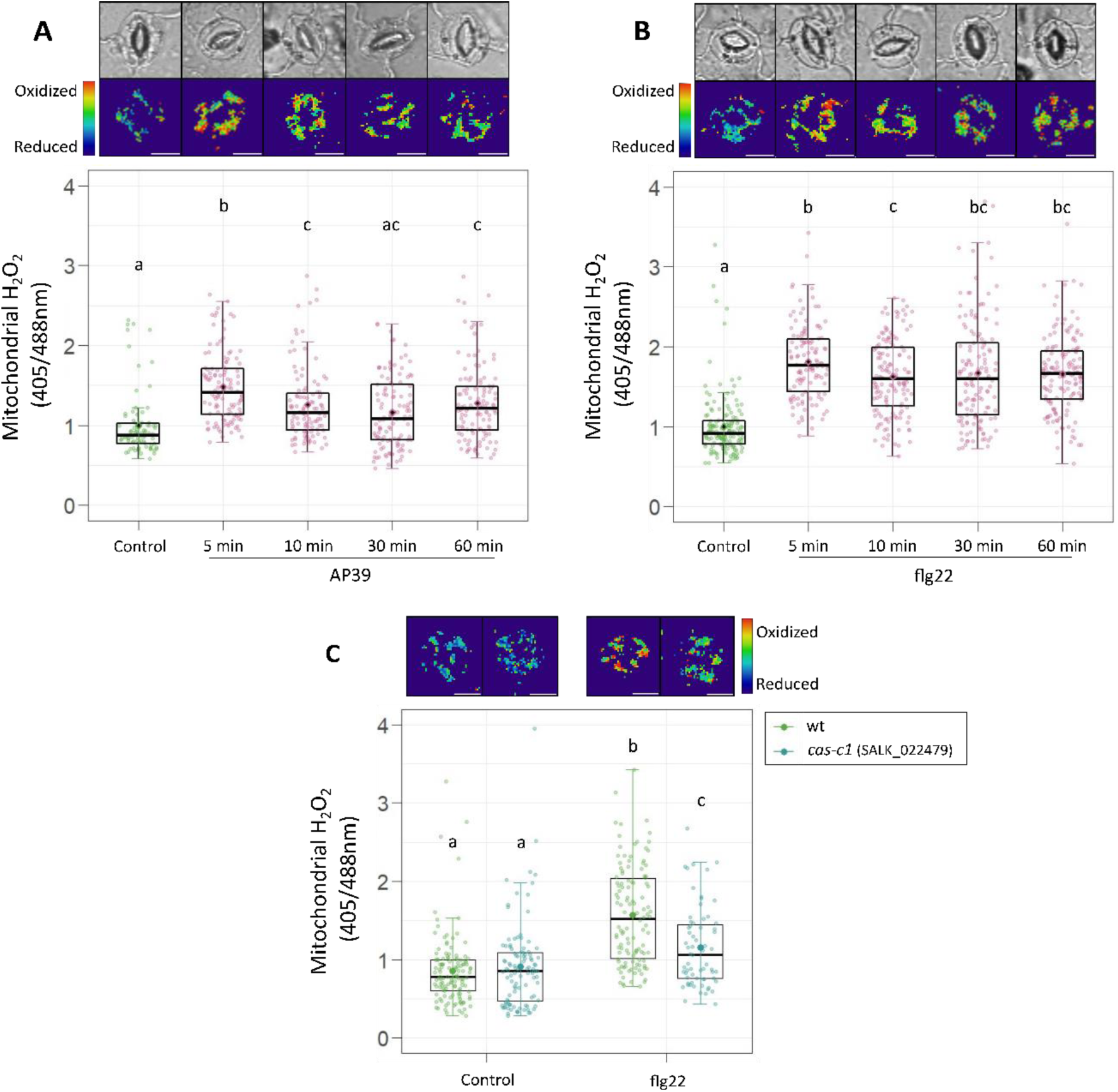
AP3G and flg22 modulates mitochondrial redox state in guard cells. Epidermal strips from *Arabidopsis thaliana* wild-type (wt) or *cas-c1* (SALK_022479) leaves expressing the mt-roGFP2-Orp1 biosensor for mitochondrial H_2_O_2_ detection were incubated in opening buffer (5 mM MES, 50 mM KCl, pH = 6.1) for 7 hr and then treated with: 100 nM AP39 or 1 µM flg22 for 5, 10, 30 and 60 min (A, B), or 5 min with flg22 (C). Guard cells fluorescence was detected by confocal microscopy, and 405/488 nm ratios were calculated for each guard cell, normalized to the control of wt and represented as individual points. Boxplots are bound by the 25^th^ to 75^th^ percentile. The line in the middle is the median, the darker point is the mean and whiskers span 10^th^ to 90^th^ percentile. Black points represent the outliers. Data is from at least three independent replicates for each experiment. Different letters indicate statistical differences among treatments (Tukey’s method, confidence level = 0.95). Ratiometric images are false-colored and represent the mean of different treatments. Scale bars: 10 μm.

Furthermore, the mitochondrial *E*_GSH_ was monitored using the mt-Grx1-roGFP2 sensor, which exhibited rapid oxidation after AP39 treatment (Supplementary Fig. S1A). These results indicate an early modulation of redox status driven by mtH_2_S in guard cells. These oxidation dynamics resemble those previously shown for the cytosolic roGFP2-Orp1 and Grx1-roGFP2 biosensors in response to the mtH_2_S donor (Pantaleno et al., 2023).

We recently reported that mitochondrial H_2_S source (CAS-C1) is required for flg22-induced cytosolic H_2_O_2_ increase in guard cells (Pantaleno et al., 2025). To underpin the physiological relevance of endogenous mtH_2_S in the control of mitochondrial redox state during stomatal immunity, we analyzed the mt-roGFP2-Orp1 response to flg22 in wt and *cas-c1* plants. First, we evaluated the mitochondrial H_2_O_2_ dynamics in guard cells from wt epidermal peels treated with flg22 at different time points. Figure 1 shows that mt-roGFP2-Orp1 is rapidly oxidized after 5 minutes of PAMP exposure and remains oxidized for at least 60 minutes (Fig. 1B). While flg22-induced mitochondrial oxidation has been reported in whole leaf tissue (Nietzel et al., 2019; Arnaud et al., 2023a), these data resolve this response at the guard cell level. We then analyzed flg22-triggered mitochondrial H_2_O_2_ increase in guard cells from *cas-c1* plants (SALK_022479). Fluorescence measurements indicate that flg22-induced sensor oxidation was significantly attenuated, although not abolished, in the *cas-c1* background as compared with wt (Fig. 1C). The observed increase of mitochondrial H_2_O_2_ was also impaired in an independent insertional *cas-c1* line (SALK_068045C), pointing to an affected mitochondrial redox response in the mutant (Supplementary Fig. S1B). Together, these findings imply that mitochondrial redox dynamics participate in the signalling events that control stomatal immunity, while also suggesting a regulatory role for mtH_2_S/CAS-C1 in this response.

### Functional mitochondria are required for stomatal closure and the increase of mtH_2_O_2_ levels

Mitochondria have long been implicated in biotic stress responses in plants (Colombatti et al., 2014; Wang et al., 2022), however, the role of mitochondrial activity in stomatal closure upon PAMP recognition has received little attention. We have shown that AP39 increases Δψm in guard cells, while blocking the mETC with different inhibitors, affects both flg22 and AP39-triggered stomatal closure (Pantaleno et al., 2023; Pantaleno et al., 2025). Therefore, we addressed the relevance of Δψm for stomatal closure and mitochondrial H_2_O_2_ increase in guard cells, in response to these stimuli. Following that aim, we measured the stomatal closure response and the mt-roGFP2-Orp1 fluorescence in wt epidermal strips treated with flg22 or AP39, in the presence or absence of the protonophore 2,4-dinitrophenol (DNP), which dissipates the proton gradient across the inner mitochondrial membrane, inducing membrane depolarization in a fraction of mitochondrial population (Ma et al., 2021). Figure 2 shows that while flg22 and AP39 treatments induced stomatal closure and mt-roGFP2-Orp1 oxidation, incubation with the uncoupler impaired both responses (Fig. 2). These results indicate that mitochondrial electrochemical potential and thereby normal mETC activity, is needed for mtH_2_S-and flg22-mediated mitochondrial redox regulation and stomatal closure induction.

**Figure 2.**
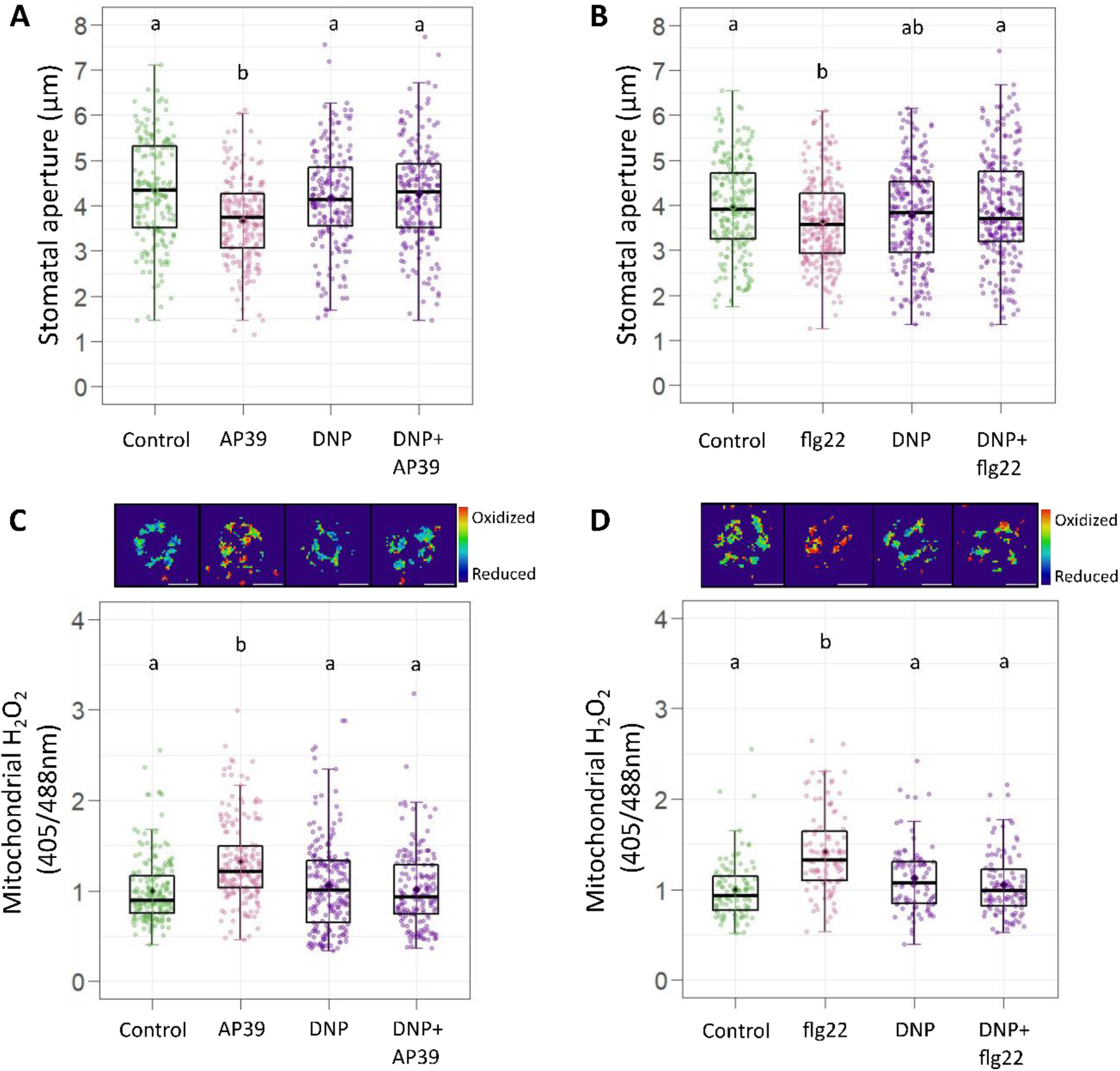
mETC activity is required for mtH₂S and flg22-induced mtH₂O₂ increase and stomatal closure. *Arabidopsis thaliana* wild-type (wt) epidermal peels were isolated and subsequently floated in opening buffer for 3 hr in the light. Then, for stomatal aperture measurements, they were treated for 90 min with 100 nM AP39, 1 µM flg22, 2 hr with 50 µM 2,4-dinitrophenol (DNP) or pre-incubated 30 min with DNP followed by a 90 min incubation with AP39 or flg22 as indicated (A,B). To monitor mitochondrial H_2_O_2_, epidermal strips from wt plants expressing mt-roGFP2-Orp1 were incubated in opening buffer for 7 hr and then treated with: 100 nM AP39 or 1 µM flg22 for 5 min, 50 µM DNP 1 hr, or preincubated with the uncoupler and then treated with AP39 or flg22 (C,D). Stomatal aperture (µm) and normalized 405/488 nm ratios are represented as points in the boxplots, where the box is bound by the 25^th^ to 75^th^ percentile. The line in the middle is the median, the darker point is the mean and whiskers span 10^th^ to 90^th^ percentile. Data from each experiment is from at least three independent replicates. Different letters indicate statistical differences among treatments (Tukey’s method, confidence level = 0.95). Ratiometric images are false-colored and represent the mean of different treatments. Scale bars: 10 μm.

### CYTc participates in stomatal immunity

Mitochondrial CYTc regulates plant responses to different environmental stresses (Racca et al., 2018; Racca et al., 2022; Canal et al., 2024; Coronel et al., 2025). Given the requirement of functional mitochondria for AP39 and flg22-elicited stomatal closure, and the fundamental role of CYTc in the mETC activity, we assessed the impact of this protein in stomatal movement.

Previously, we reported that *cytc* knock-down double mutants were unable to induce stomatal closure in response to AP39 (Pantaleno et al., 2023). To investigate the participation of CYTc in the stomatal response to PAMPs, we challenged epidermal strips from two independent *cytc* lines (*1*b*2*a and *1*b*2*b) with flg22. Stomatal pore width measurements showed that both mutants were impaired to induce stomatal closure upon flg22 treatment, evidencing the participation of CYTc in this early immune response (Fig. 3A). Further, to evaluate if CYTc is specifically involved in the stomatal closure triggered by biotic stimuli, we treated isolated epidermal peels with ABA or two other synthetic PAMPs from bacteria or fungi, elf18 or chitin, respectively. As shown in Figure 3B, ABA-induced stomatal closure was unaffected in mutant plants. In contrast, the stomatal response to elf18 and chitin was impaired in both *1*b*2*a and *1*b*2*b lines (Fig. 3B). To rule out the possibility that *cytc* mutants exhibit broadly impaired stomatal movement regulation, we evaluated light-induced stomatal opening over time in wt and the *cytc* lines, and we observed a similar response between all genotypes (Fig. 3C). These results underscore the CYTc requirement for PAMP-elicited stomatal immunity and highlight the relevance of mitochondrial activity in the stomatal response to biotic stress.

**Figure 3.**
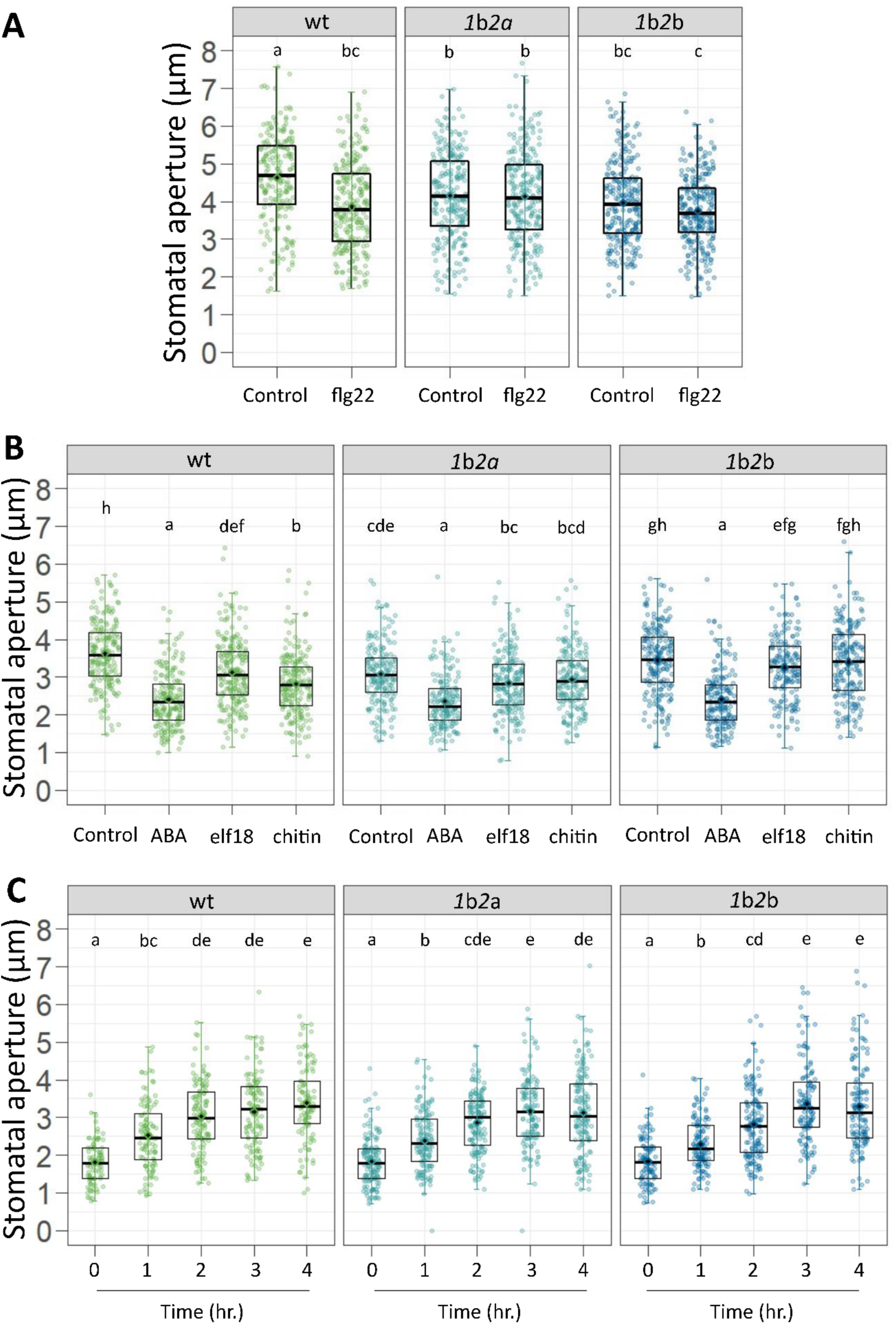
CYTc is required for PAMP-triggered stomatal closure. Isolated epidermal peels were excised from the abaxial side of leaves from 5-week-old *Arabidopsis thaliana* wild-type (wt) or *cytc* double mutants (*1*b*2*a and *1*b*2*b). For stomatal opening assays strips were subsequently floated in opening buffer and immediately imaged (0 hr) or after 1, 2, 3 or 4 h of incubation in the light (C). To assess stomatal closure, peels were incubated for 3 hr in the light and treated for 90 min with: 1 µM flg22, 1 µM elf18, 1mg/ml chitin or 20 µM abscisic acid (ABA) (A,B). Stomatal aperture absolute values in microns (µm) are represented as points in the boxplots, where the box is bound by the 25^th^ to 75^th^ percentile. The line in the middle is the median, the darker point is the mean and whiskers span 10^th^ to 90^th^ percentile. Data was taken from at least three independent biological replicates for each experiment. Different letters indicate statistical differences among treatments (Tukey’s method, confidence level = 0.95).

The function of CYTc as an electron transporter in the mETC has a fundamental impact on the establishment of the Δψm needed for ATP synthesis (Welchen and Gonzalez, 2016). Since mitochondrial depolarization affected the stomatal response (Fig. 2), we analyzed the CYTc-dependent Δψm modulation in guard cells incubated with either AP39 or flg22. With that aim we loaded epidermal strips from wt or *cytc* plants with the fluorescent probe TMRM, and then we treated them with the mtH_2_S donor, the bacterial PAMP or DNP as a control. Guard cell-specific fluorescence intensity quantification shows that both AP39 and flg22 increased mitochondrial polarization in wt guard cells. This response was not observed in either *cytc* line (Fig. 4A-C), suggesting that CYTc-dependent mitochondrial hyperpolarization participates in the signalling events that drive stomatal immunity.

**Figure 4.**
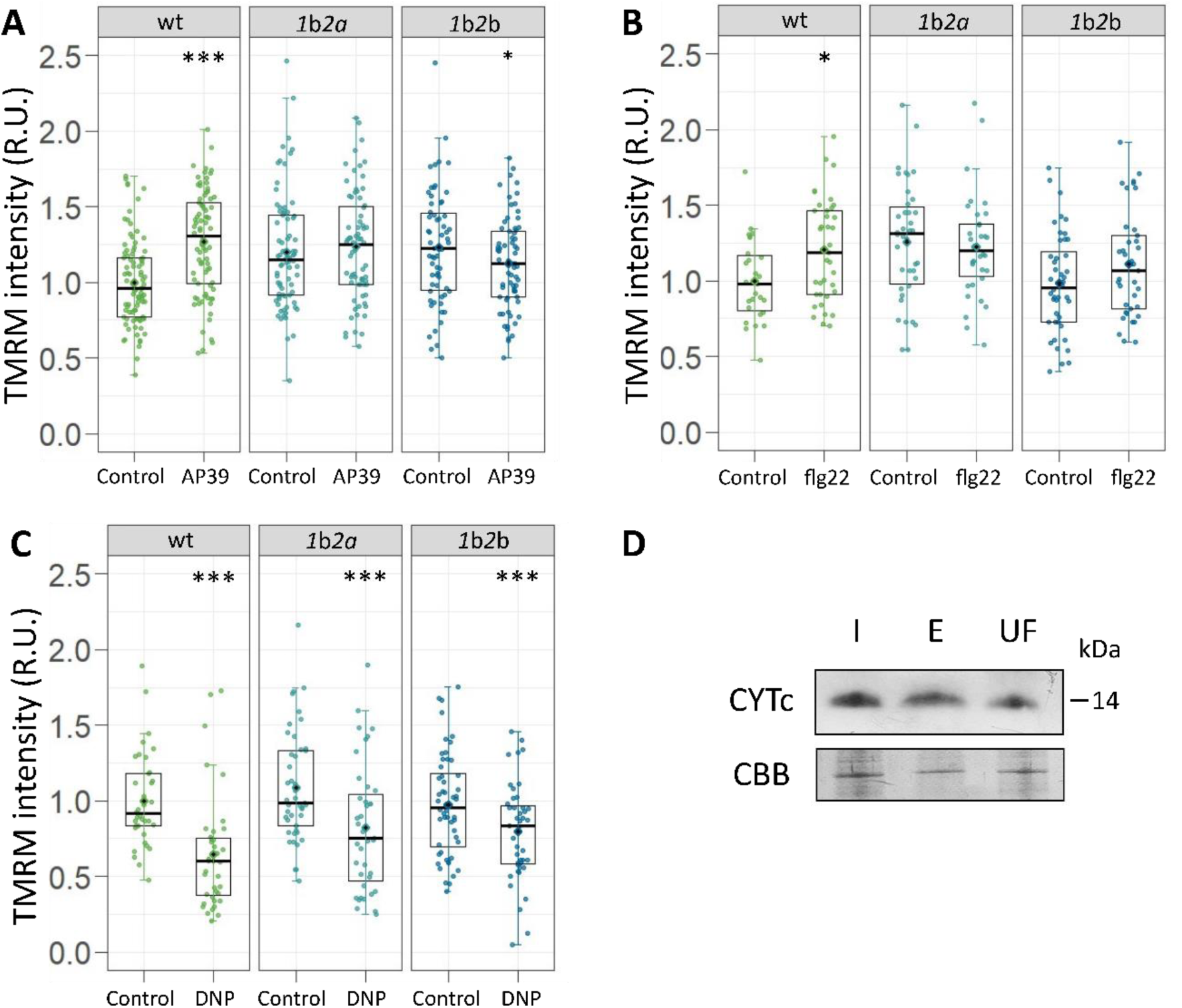
CYTc is persulfidated *in vivo* and is required for flg22 and AP3G-driven mitochondrial hyperpolarization. Isolated epidermal peels were excised from the abaxial side of leaves from 5-week-old *Arabidopsis thaliana* wild-type (wt) or *cytc* double mutants (*1*b*2*a and *1*b*2*b) (A,B,C). For mitochondrial inner membrane potential determination, epidermal strips were incubated in opening buffer supplemented with 500 nM Tetramethylrhodamine methyl ester (TMRM) in the dark and then treated with 100 nM AP39 (A), 1 µM flg22 (B) for 5 min or with 50 µM DNP for 1 hr as a control of the experimental setup (C). TMRM intensity was calculated for each guard cell and normalized to the wt control. TMRM relative units (R.U.) of fluorescence are represented as points in the boxplots, where the box is bound by the 25^th^ to 75^th^ percentile. The line in the middle is the median, the darker point is the mean and whiskers span 10^th^ to 90^th^ percentile. Data was taken from at least three independent biological replicates for each experiment. Asterisks indicate statistical differences between treatments within each genotype (Dunnett’s method, confidence level = 0.95). (D) Western blot analysis of persulfidated proteins purified by dimedone-switch method (I: input, E: eluate, and UF: unbound fraction). An anti-CYTc (CYTc) antibody was used to detect CYTc, and Coomassie Brilliant Blue (CBB) staining was used as a loading control. Similar results were obtained in independent biological replicates (Supplementary Figure S2).

Since H_2_S-mediated persulfidation was reported to regulate protein activity, we studied CYTc persulfidation *in vivo*. Persulfidated proteins were enriched by the dimedone-switch method (Aroca et al., 2022), and CYTc was subsequently identified by Western blot. Results presented in Figure 4 indicate that CYTc is detected in both the eluted (persulfidated) and unbound (non-persulfidated) protein fractions (Fig. 4D, Supplementary Fig. S3). These findings provide *in vivo* validation of CYTc as a target of H_2_S-driven posttranslational modification.

### CYTc is required for mtH_2_S and flg22-dependent redox regulation in guard cell cytosol

Stomatal immune response requires an increase in intracellular H_2_O_2_, which has been mainly attributed to the well-established PAMP-triggered apoplastic ROS production driven by the NADPH oxidase Respiratory Burst Oxidase Homolog D (RBOHD) (Kadota et al., 2014; Li et al., 2014; Qi et al., 2017). Then, to analyze the participation of CYTc on this signalling pathway, we first measured flg22-induced apoplastic ROS production in Arabidopsis leaf discs from wt or *cytc* plants and we observed a similar apoplastic ROS burst in both lines, suggesting that CYTc is not involved in RBOHD-driven apoplastic ROS production (Supplementary Fig. S3).

In previous work, we showed that both flg22 and AP39 consistently oxidise the cytosolic roGFP2-Orp1 sensor in guard cells (Pantaleno et al., 2023; Pantaleno et al., 2025; Scuffi et al., 2025). Therefore, we evaluated guard-cell cytosolic redox dynamics in isolated epidermal peels from *cytc* plants expressing the sensor roGFP2-Orp1, in response to flg22 or AP39. At resting conditions, guard cells from *cytc* exhibited higher roGFP2-Orp1 oxidation levels than wt (Fig. 5). In addition, both AP39 and flg22 generated an H_2_O_2_ increase in wt plants, a response that was not observed in the *cytc* double mutant (Fig. 5A,B). These results highlight the role of mitochondrial CYTc in modulating the guard-cell cytosolic redox state upon mtH_2_S and flg22 exposure.

**Figure 5.**
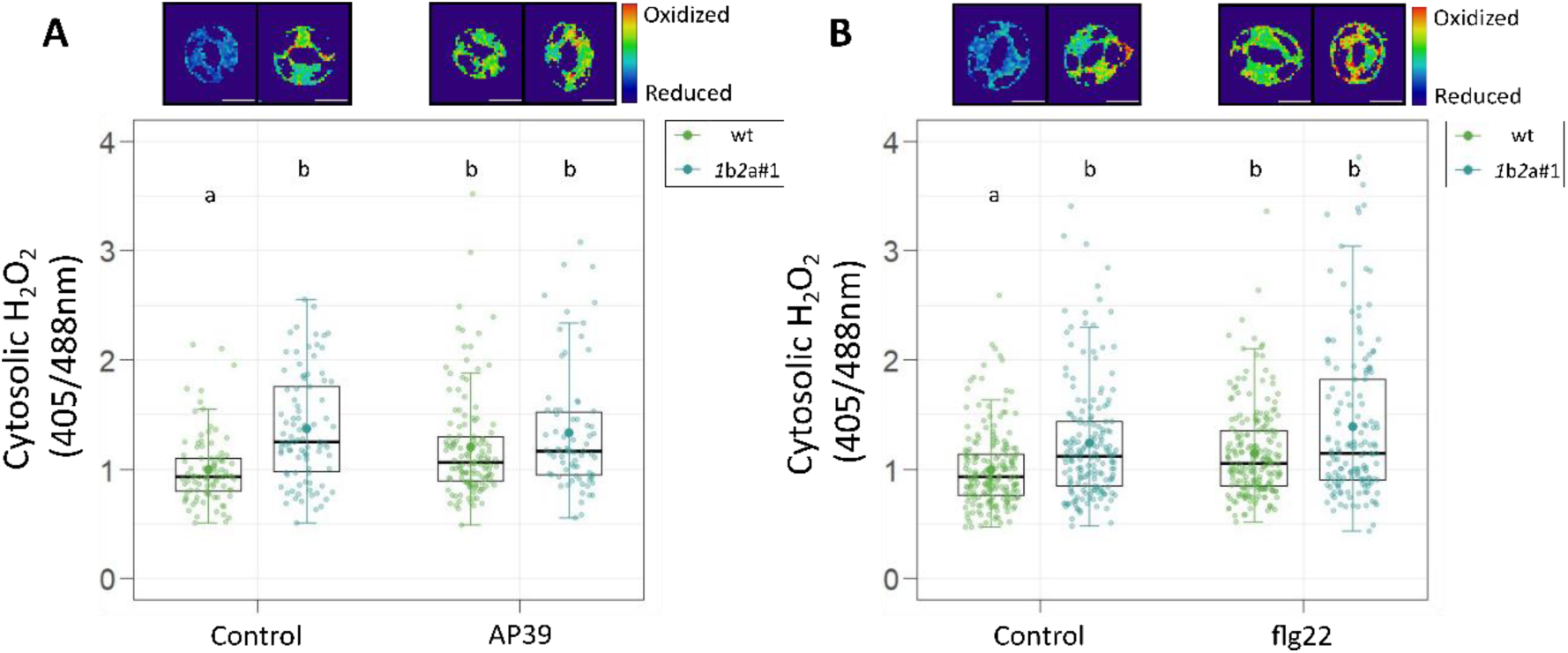
CYTc is needed for flg22 and AP3G-induced H_2_O_2_ increase in guard cells. Epidermal strips from *Arabidopsis thaliana* wild-type (wt) or *cytc 1*b*2*a double mutant expressing the roGFP2-Orp1 biosensor for cytosolic H_2_O_2_ detection were incubated in opening buffer for 7 hr and then treated with: 100 nM AP39 or 1 µM flg22 for 60 min as previously described (Pantaleno et al., 2023; Pantaleno et al., 2024). Guard cells fluorescence was detected by confocal microscopy, and 405/488 nm ratios were calculated for each guard cell, normalized to the control of wt and represented as individual points. Boxplots are bound by the 25^th^ to 75^th^ percentile. The line in the middle is the median, the darker point is the mean and whiskers span 10^th^ to 90^th^ percentile. Black points represent the outliers. Data is from at least three independent replicates for each experiment. Different letters indicate statistical differences among treatments (Tukey’s method, confidence level = 0.95). Ratiometric images are false-colored and represent the mean of different treatments. Scale bars: 10 μm.

### Plant mutants lacking CYTc are more susceptible to *Pseudomonas* infection

To further investigate the regulatory role of CYTc on the stomatal response during PTI, we assessed the impact of CYTc-mediated stomatal closure on whole plant immune response. First, we assayed the requirement of CYTc for the stomatal response to bacteria by treating epidermal strips from *cytc* plants with a suspension of *Pseudomonas syringae* pv. *tomato* DC3000 (*Pst*. DC3000) or the type III secretion system mutant *hrcC^-^*(*Pst*. *hrcC^-^*), which is unable to deliver effectors to counteract PTI (Zipfel et al., 2004). Incubation with *Pst*. DC3000 triggered stomatal closure in both wt and *cytc*, while the mutant presented an impaired stomatal response to *Pst*. *hrcC^-^,* providing further evidence for the implication of CYTc in stomatal immunity response elicited by PAMPs (Fig. 6A).

**Figure 6:**
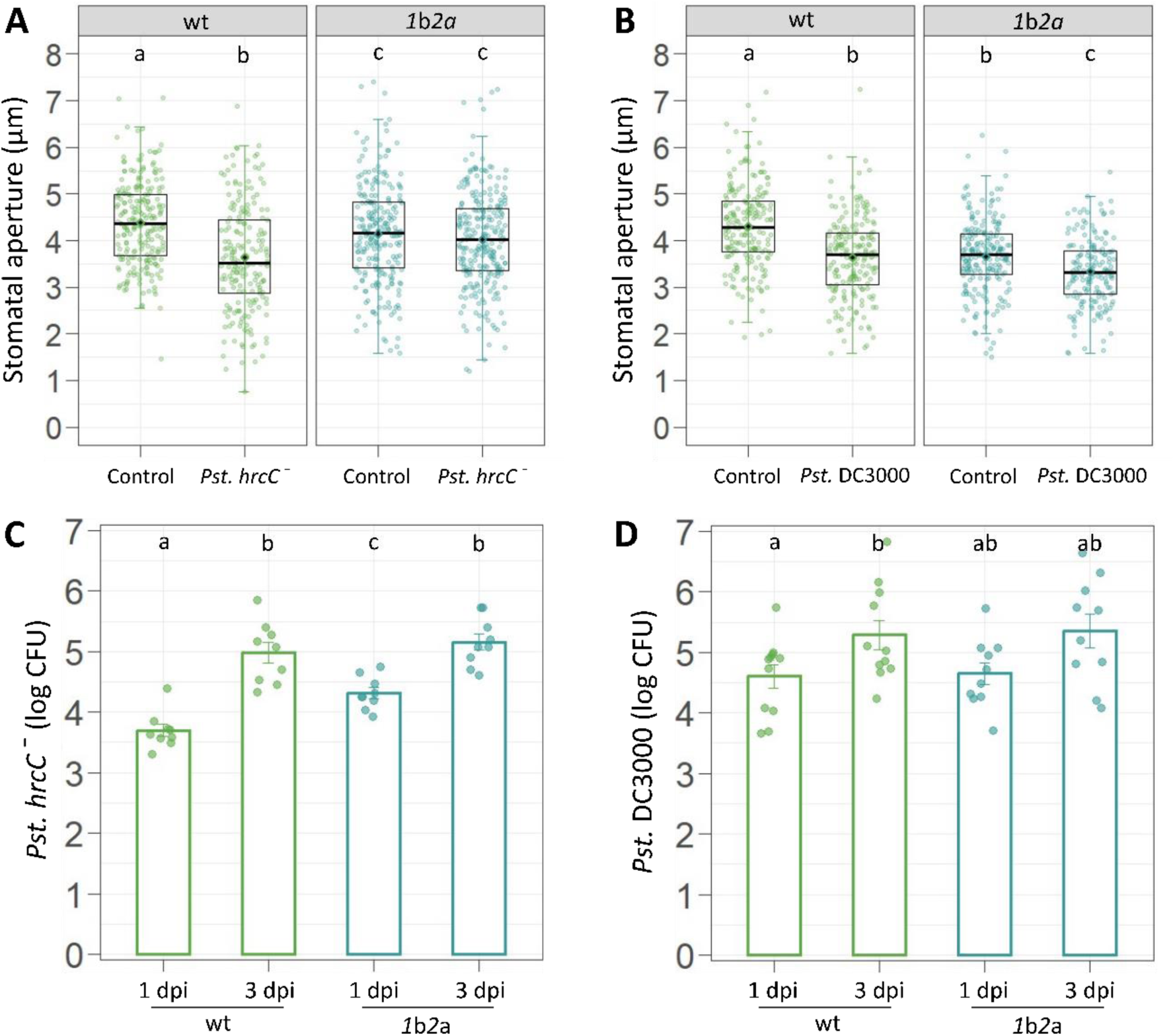
***cytc* mutants are more susceptible to *Pst. hrcC^-^* infection.** Stomatal aperture was measured on isolated epidermal strips from *Arabidopsis thaliana* wild-type (wt) or *cytc* double mutant (*1*b*2*a) leaves. Peels were floated in opening buffer for 3 h in light and treated for 60 min with a *Pseudomonas syringae* pv. *tomato* DC3000 wt (*Pst*. DC3000) or *hrcC^-^* (*Pst. hrcC^-^*) suspension (OD = 0.1 (λ = 600 nm)). Stomatal pore size absolute values in microns (µm) are represented as points in the boxplots, where the box is bound by the 25^th^ to 75^th^ percentile. The line in the middle is the median, the darker point is the mean and whiskers span 10^th^ to 90^th^ percentile. For whole plant immunity assessment, wt and *cytc* plants were surface-inoculated by spray with a *Pst*. DC3000 or *Pst. hrcC^-^* suspension (OD = 0.2 (λ = 600 nm)), and bacterial growth was determined in leaf discs at 1- or 3-days postinoculation (dpi). The number of colony-forming units (CFU)/cm^2^ of leaf extracts was determined for each plant and showed as individual points. Bars represent the mean bacterial growth values, error bars are the standard error. Different letters indicate statistical differences among conditions (Tukey’s method, confidence level = 0.95).

To address the role of CYTc in whole-plant immunity, we infected plants with *Pst*. DC3000 or *Pst*. *hrcC^-^*by spray-based inoculation, a method that allows analysis of the functional impact of stomatal closure on pathogen invasion. At 1 day post-infection (dpi), *cytc* plants inoculated with *Pst*. *hrcC^-^*presented significantly higher bacterial growth with respect to wt, whereas this difference was not observed in plants infected with the *Pst.* DC3000 strain (Fig. 6C,D). Notably, these results correlate with the stomatal responses to bacterial challenge. After 3 dpi, there were no differences between the mutants and wt plants. These findings indicate CYTc participation in eliciting stomatal immunity during early infection stages, ultimately impacting whole-plant defence.

Together, these findings support a role for mtH_2_S in the CYTc-dependent modulation of guard cell redox state and stomatal closure upon PAMP perception, with a concomitant impact on plant immunity.

## Discussion

The present study provides new evidence for a central role of mitochondrial activity in regulating stomatal closure. These findings reveal concerted action between guard cell mtROS and mETC activity in activating flg22-triggered stomatal immunity, a process that requires the mitochondrial H_2_S source CAS-C1 and the mETC component CYTc.

Stomatal immunity induced by PAMPs has been widely studied over the past decades. As a result, we now have a relatively detailed understanding of the signalling network that commands this response. However, breakthroughs in emerging technologies, including biosensors that enable for specific, high-resolution, and *in vivo* monitoring of target ions and molecules at the subcellular level, continue revealing novel components and regulatory layers within guard cell signaling pathways.

PAMP perception triggers transient ROS bursts in different subcellular compartments (Nietzel et al., 2019; Arnaud et al., 2023a). Although the role of mtROS role in stomatal immunity is not well described, it was recently reported that perception of flg22 elicits an increase in mtROS in different plant tissues (Nietzel et al., 2019; Arnaud et al., 2023a; Yang et al., 2022; Yang et al., 2025), while the pharmacologic scavenging of mtROS impaired flg22-induced stomatal closure (Lu et al., 2026).

Here we found that DNP-mediated mitochondrial depolarization impaired stomatal closure and the early mitochondrial H_2_O_2_ increase in guard cells driven by AP39 and flg22, showing that an intact Δψm is needed for both stomatal movement and mitochondrial redox state modulation (Fig. 2). These observations suggest that mETC-dependent mtROS increase might have a prominent role in the activation of stomatal immunity. Furthermore, the flg22-elicited H_2_O_2_ increase in mitochondrial matrix was also attenuated in the guard cells from *cas-c1* plants (Fig 1C). Consistent with the PAMP-induced mitochondrial H_2_O_2_ response, we previously showed that *cas-c1* guard cells also exhibit a diminished cytosolic H_2_O_2_ increase upon flg22 exposure. Altogether, these data point towards a link between endogenous mtH_2_S and the mitochondrial and cytosolic redox regulation during stomatal immunity (Pantaleno et al., 2023; Pantaleno et al., 2025).

The central function of CYTc in mitochondrial bioenergetics is evidenced in the *cytc* plants phenotype, which exhibit decreased ATP contents (Racca et al., 2018; Racca et al., 2022; Canal et al., 2024). Previous data shows that *cytc* roots cells exhibited lower Δψm than wt plants (Canal et al., 2024), a difference that was not evident in guard cells (Fig. 4A,B,C). This apparent discrepancy can be attributed to i) a cell-specific regulation of CYTc activity; ii) the resolution of the readout or iii) differences related to the experimental systems and methodological designs.

Recently we reported that CYTc is required for the induction of stomatal closure in response to mtH_2_S (Pantaleno et al., 2023). In the present work we show that *cytc* mutants have compromised the PAMP-elicited stomatal closure, and the flg22 and mtH_2_S-dependent mitochondrial hyperpolarization (Fig. 3 and 4), underscoring a potential role for mitochondrial activity in the stomatal response regulation. In addition, given that independent proteomic datasets identified CYTc as a target of persulfidation (García-Calderón et al., 2023; Jurado-Flores et al., 2023), we studied this process *in vivo*. The results reveal that CYTc undergoes an H_2_S-dependent posttranslational modification in plants, validating the persulfidation of this mitochondrial protein (Fig. 4D, Supplementary Fig. S2).

CYTc deficiency impaired stomatal closure in response to three structurally unrelated PAMPs (flg22, elf18 and chitin) but did not affect ABA-induced closure or light-induced opening (Fig. 3). These results suggest that guard cells with reduced CYTc remain functional, but the lack of mitochondrial input specifically affects PAMP-induced stomatal closure. Dissipating Δψm with DNP blocked stomatal closure (Fig. 2), whereas the milder bioenergetic defect of the *cytc* knock-down lines affected only the PAMP response. This points to a Δψm-dependent mitochondrial signal, such as mtROS, rather than ATP supply alone as the limiting factor in PAMP-triggered closure. An intact ABA response was not necessarily expected, because CYTc levels modulate ABA-dependent seed germination through ABI4 (Racca et al., 2022). The connection between CYTc and ABA signalling therefore seems to depend on the process and tissue involved. The interplay between mitochondria and stomatal immunity was recently addressed in a work focused on the role of mitochondrial fusion in flg22-induced stomatal closure (Lu et al., 2026). The authors observed a decrease in Δψm and ATP content in guard cells of seedlings incubated for 30 minutes with flg22. In contrast, we detected a CYTc-dependent increase in TMRM fluorescence after 10 minutes (Fig. 4). This transient mitochondrial dysfunction, which may arise as a downstream effect of immune stress, was then restored by MIRO1-mediated mitochondrial fusion. Consequently, *miro1* plants failed to recover mitochondrial function and presented an impaired flg22-triggered stomatal closure and increased susceptibility to bacterial infection (Lu et al., 2026), highlighting the relevance of functional mitochondria for stomatal immunity. Rather than being contradictory, our observations are compatible with a biphasic response in which early, CYTc-dependent hyperpolarization and associated mtROS production precede mitochondrial dysfunction, which is later repaired by MIRO1-mediated fusion.

PAMP perception produces an increase in cellular ROS levels, which is mainly attributed to an RBOHD-driven apoplastic ROS burst (Kadota et al., 2014; Li et al., 2014). Dissection of intracellular redox dynamics in guard cells indicated that the cytosolic H_2_O_2_ increase, triggered by either AP39 or flg22, is absent in the *cytc* double mutant, which also presented a higher oxidized redox state under basal conditions (Fig. 5). These results implicate a cellular redox imbalance resembling the altered H_2_O_2_ dynamics reported for guard cell cytosol from *cas-c1* plants (Pantaleno et al., 2025). Interestingly, a previous finding indicated that *cytc* seedlings exhibited lower ROS levels under resting state (Coronel et al., 2025). This difference can also be associated with the different experimental conditions, including the specificity of the detection system, the plants developmental stage and the distinctive properties of the tissue/cells analyzed in each case.

The observation that apoplastic H_2_O_2_ production upon flg22 treatment is similar between wt and *cytc* plants (Supplementary Fig. S3), indicates an apparent RBOHD-independent modulation of cytosolic H_2_O_2_. Consistent with these results, several studies have hypothesized that the observed pathogen-induced increases in cytosolic and mitochondrial ROS are not related to the RBOHD-mediated apoplastic ROS burst (Pei et al., 2022; Arnaud et al., 2023b; Yang et al., 2025). Higher mtROS levels triggered upon *Phytophthora parasitica* exposure remained unaffected in the presence of inhibitors of RBOH activity (Yang et al., 2025), while flg22-triggered mtROS increase was similar between wt plants and *rbohD* mutants (Yang et al., 2022). Moreover, it was recently shown that mtROS are transferred to the endoplasmic reticulum (ER), suggesting that mitochondria may act as a cellular ROS source to induce immune responses (Yang et al., 2025). However, the interplay between mtROS and cytosolic redox state in guard cells during stomatal immunity requires further investigation.

The *cytc* mutants are also compromised in their immune response at the whole-plant level. Following spray inoculation with *Pst. hrcC⁻*, the *cytc* plants displayed an early increased susceptibility within 1 dpi, consistent with its impaired stomatal closure response to the bacterial suspension (Fig. 6A,C). Conversely, both stomatal immunity and bacterial resistance were comparable between wt and *cytc* plants upon *Pst*. DC3000 inoculation (Fig. 6B,D).

Given that virulent *Pst.* DC3000 suppresses PTI and stomatal immunity through type III effectors and the phytotoxin coronatine, whereas *Pst. hrcC*_−_ cannot deliver effectors and is therefore restricted mainly by PTI, the presented data point to CYTc as a component of PTI-associated stomatal immunity. These results highlight the critical contribution of PAMP-triggered stomatal closure to plant defense, while suggest that the early susceptibility of *cytc* can be related to defective stomatal immunity. These findings contribute to understanding the relevance of mETC activity on whole-plant immune response.

Virulent bacteria activate hormone-dependent pathways, which may bypass the PAMP-specific branch defective in *cytc*. Previous findings indicate that *cytc* plants show enhanced SnRK1 signalling (Coronel et al., 2025) and lower ATP levels (Racca et al., 2018) and either could affect post-invasive defense. However, the susceptibility phenotype was transient and appeared only in *Pst. hrcC*⁻, suggesting a mainly pre-invasive, stomatal contribution. The results show that the signalling molecule mtH_2_S regulates Δψm and mitochondrial redox status in guard cells, in a CYTc-dependent manner, to trigger the stomatal immune response. Overall, CYTc is required for stomatal closure induced by PAMPs but not by ABA, and for the changes in guard cell Δψm and cytosolic redox state elicited by both flg22 and mtH_2_S. Together with the attenuated flg22-induced mitochondrial oxidation in *cas-c1* and the *in vivo* persulfidation of CYTc, these results support a model in which mitochondrial H_2_S signalling converges on CYTc and modulates guard cell redox dynamics during stomatal immunity.

In addition, we observed that CYTc is required for PAMP-induced stomatal closure, providing new evidence on the participation of mitochondrial bioenergetics in plant immunity. Altogether, these findings show that mitochondrial activity and mtROS modulation in guard cells make a fundamental contribution to the signalling network that orchestrates stomatal closure, positioning mitochondria as a key player in plant immunity activation.

## Accession numbers

*CAS-C1* (At3g61440), *CYTC-1* (At1g22840), *CYTC-2* (At4g10040).

## Supporting information

Supplementary Material Pantaleno

## Acknowledgements

This work was financially supported by Universidad Nacional de Mar del Plata (UNMDP) and Consejo Nacional de Investigaciones Científicas y Técnicas (CONICET). PICT 2021-GRF1-092 (to DS), PICT 2021-0189 (to CGM), postdoctoral fellowship from CONICET (to RP).

## Author contributions

R.P. performed most of the experiments, analyzed the data, and wrote the article; D.S. performed dimedone-switch method, analyzed the data, and helped with the writing of the article; F.C. did the CYTc inmunodetection, E.W. participated in the discussion and helped with the writing of the article, and C.G.M. conceived the project and wrote the article.

## Supplementary material

**Supplementary Figure S1: AP3G and flg22 modulates mitochondrial redox state in guard cells.** Epidermal strips from *Arabidopsis thaliana* wild-type (wt) leaves expressing the mt-Grx1-roGFP2 biosensor for mitochondrial *E*_GSH_ detection (A) or wt and *cas-c1* (SALK_ 068045C) expressing mt-roGFP2-Orp1 to monitor mitochondrial H_2_O_2_ (B), were incubated in opening buffer (5 mM MES, 50 mM KCl, pH = 6.1) for 7 hr and then treated with: 100 nM AP39 for 5, 10, 30 and 60 min (A) or 1 µM flg22 for 5 min (B). Guard cells fluorescence was detected by confocal microscopy, and 405/488 nm ratios were calculated for each guard cell, normalized to the control of wt and represented as individual points. Boxplots are bound by the 25^th^ to 75^th^ percentile. The line in the middle is the median, the darker point is the mean and whiskers span 10^th^ to 90^th^ percentile. Black points represent the outliers. Data is from at least three independent replicates for each experiment. Different letters indicate statistical differences among treatments (Tukey’s method, confidence level = 0.95).

**Supplementary Figure S2: CYTc is persulfidated *in vivo*.** Western blot analysis of persulfidated proteins purified by dimedone switch method (I: input, E: eluate, and UF: unbound fraction). An anti-CYTc (CYTc) antibody was used to detect CYTc, and Coomassie Brilliant Blue (CBB) staining was used as a loading control.

**Supplementary Figure S3: Apoplastic ROS production in *cytc* mutants.** Leaf discs from 5-week-old *Arabidopsis thaliana* wild-type (wt) or *cytc* double mutant (*1*b*2*a) plants were floated in deionized water overnight. The following day, they were placed on 96-well white plate containing a solution with 20 mM luminol and 0.02 mg/mL horseradish peroxidase. ROS production was triggered with 100 nM flg22 and luminescence was measured with a luminometer. Apoplastic ROS production is expressed as the relative light units (RLU) detected (A), or the area under the curve (AUC) for each leaf disc (B). In (A) the points represent the mean RLU over time and the error bars are the standard error. In (B) the AUC are represented as points in the boxplots, where the box is bound by the 25^th^ to 75^th^ percentile. The line in the middle is the median, the darker point is the mean and whiskers span 10^th^ to 90^th^ percentile. Black points represent measurements outside those margins. Data was taken from at least three independent experiments. Different letters indicate statistical differences among treatments (Tukey’s method, confidence level = 0.95).

## References

Arnaud D, Deeks MJ, Smirnoff N (2023a) Organelle-targeted biosensors reveal distinct oxidative events during pattern-triggered immune responses. Plant Physiology 191: 2551–2569

Arnaud D, Deeks MJ, Smirnoff N (2023b) RBOHF activates stomatal immunity by modulating both reactive oxygen species and apoplastic pH dynamics in Arabidopsis. The Plant Journal 116: 404–415

Aroca A, Gotor C, Bassham DC, Romero LC (2020) Hydrogen Sulfide: From a Toxic Molecule to a Key Molecule of Cell Life. Antioxidants 9: 621

Aroca A, Jurado-Flores A, Filipovic MR, Gotor C, Romero LC (2022) Detection of protein persulfidation in plants by the dimedone switch method. Methods in Enzymology. Elsevier, pp 385–402

Aroca A, Yruela I, Gotor C, Bassham DC (2021) Persulfidation of ATG18a regulates autophagy under ER stress in *Arabidopsis*. Proc Natl Acad Sci USA 118: e2023604118

Benchoam D, Cuevasanta E, Möller M, Alvarez B (2019) Hydrogen Sulfide and Persulfides Oxidation by Biologically Relevant Oxidizing Species. Antioxidants 8: 48

Bestwick CS, Brown IR, Bennett MHR, Mansfield JW (1997) Localization of Hydrogen Peroxide Accumulation during the Hypersensitive Reaction of Lettuce Cells to Pseudomonas syringae pv phaseolicola. The Plant Cell 9: 209–221

Braun H-P (2020) The Oxidative Phosphorylation system of the mitochondria in plants. Mitochondrion 53: 66–75

Canal MV, Mansilla N, Gras DE, Ibarra A, Figueroa CM, Gonzalez DH, Welchen E (2024) Cytochrome *c* levels affect the TOR pathway to regulate growth and metabolism under energy-deficient conditions. New Phytologist 241: 2039–2058

Chen S, Jia H, Wang X, Shi C, Wang X, Ma P, Wang J, Ren M, Li J (2020) Hydrogen Sulfide Positively Regulates Abscisic Acid Signaling through Persulfidation of SnRK2.6 in Guard Cells. Molecular Plant 13: 732–744

Colombatti F, Gonzalez DH, Welchen E (2014) Plant mitochondria under pathogen attack: A sigh of relief or a last breath? Mitochondrion 19: 238–244

Coronel FP, Gras DE, Canal MV, Roldan F, Welchen E, Gonzalez DH (2025) Cytochrome *c* levels link mitochondrial function to plant growth and stress responses through changes in SnRK1 pathway activity. The Plant Journal 121: e17215

Cvetkovska M, Vanlerberghe GC (2012) Alternative oxidase modulates leaf mitochondrial concentrations of superoxide and nitric oxide: Rapid report. New Phytologist 195: 32–39

DeFalco TA, Zipfel C (2021) Molecular mechanisms of early plant pattern-triggered immune signaling. Molecular Cell 81: 3449–3467

Filipovic MR, Zivanovic J, Alvarez B, Banerjee R (2018) Chemical Biology of H _2_ S Signaling through Persulfidation. Chem Rev 118: 1253–1337

Florez-Sarasa I, Welchen E, Racca S, Gonzalez DH, Vallarino JG, Fernie AR, Ribas-Carbo M, Del-Saz NF (2021) Cytochrome c Deficiency Differentially Affects the In Vivo Mitochondrial Electron Partitioning and Primary Metabolism Depending on the Photoperiod. Plants 10: 444

Fuchs R, Kopischke M, Klapprodt C, Hause G, Meyer AJ, Schwarzländer M, Fricker MD, Lipka V (2016) Immobilized Subpopulations of Leaf Epidermal Mitochondria Mediate PENETRATION2-Dependent Pathogen Entry Control in Arabidopsis. Plant Cell 28: 130–145

García I, Castellano JM, Vioque B, Solano R, Gotor C, Romero LC (2010) Mitochondrial β-Cyanoalanine Synthase Is Essential for Root Hair Formation in Arabidopsis thaliana. The Plant Cell 22: 3268–3279

García-Calderón M, Vignane T, Filipovic MR, Ruiz MT, Romero LC, Márquez AJ, Gotor C, Aroca A (2023) Persulfidation protects from oxidative stress under nonphotorespiratory conditions in Arabidopsis. New Phytologist 238: 1431–1445

García-Mata C, Lamattina L (2010) Hydrogen sulphide, a novel gasotransmitter involved in guard cell signalling. New Phytologist 188: 977–984

Hou S, Rodrigues O, Liu Z, Shan L, He P (2024) Small holes, big impact: Stomata in plant–pathogen– climate epic trifecta. Molecular Plant 17: 26–49

Hu Y, Ding Y, Cai B, Qin X, Wu J, Yuan M, Wan S, Zhao Y, Xin X-F (2022) Bacterial effectors manipulate plant abscisic acid signaling for creation of an aqueous apoplast. Cell Host & Microbe 30: 518–529.e6

Huang S, Van Aken O, Schwarzländer M, Belt K, Millar AH (2016) The Roles of Mitochondrial Reactive Oxygen Species in Cellular Signaling and Stress Response in Plants. Plant Physiol 171: 1551–1559

Jezek M, Blatt MR (2017) The Membrane Transport System of the Guard Cell and Its Integration for Stomatal Dynamics. Plant Physiol 174: 487–519

Jin Z, Wang Z, Ma Q, Sun L, Zhang L, Liu Z, Liu D, Hao X, Pei Y (2017) Hydrogen sulfide mediates ion fluxes inducing stomatal closure in response to drought stress in Arabidopsis thaliana. Plant Soil 419: 141–152

Johansson ON, Fantozzi E, Fahlberg P, Nilsson AK, Buhot N, Tör M, Andersson MX (2014) Role of the penetration-resistance genes PEN1, PEN2 and PEN3 in the hypersensitive response and race-specific resistance in Arabidopsis thaliana. Plant Journal 79: 466–476

Jurado-Flores A, Aroca A, Romero LC, Gotor C (2023) Sulfide promotes tolerance to drought through protein persulfidation in Arabidopsis. Journal of Experimental Botany 74: 4654– 4669

Kadota Y, Sklenar J, Derbyshire P, Stransfeld L, Asai S, Ntoukakis V, Jones JD, Shirasu K, Menke F, Jones A, et al (2014) Direct Regulation of the NADPH Oxidase RBOHD by the PRR-Associated Kinase BIK1 during Plant Immunity. Molecular Cell 54: 43–55

Kim T-H, Böhmer M, Hu H, Nishimura N, Schroeder JI (2010) Guard Cell Signal Transduction Network: Advances in Understanding Abscisic Acid, CO_2_, and Ca^2+^ Signaling. Annu Rev Plant Biol 61: 561–591

Laureano-Marín AM, Aroca Á, Pérez-Pérez ME, Yruela I, Jurado-Flores A, Moreno I, Crespo JL, Romero LC, Gotor C (2020) Abscisic Acid-Triggered Persulfidation of the Cys Protease ATG4 Mediates Regulation of Autophagy by Sulfide. Plant Cell 32: 3902–3920

Lawson T, Matthews J (2020) Guard Cell Metabolism and Stomatal Function. Annu Rev Plant Biol 71: 273–302

Li L, Li M, Yu L, Zhou Z, Liang X, Liu Z, Cai G, Gao L, Zhang X, Wang Y, et al (2014) The FLS2-Associated Kinase BIK1 Directly Phosphorylates the NADPH Oxidase RbohD to Control Plant Immunity. Cell Host & Microbe 15: 329–338

Lin P-A, Chen Y, Ponce G, Acevedo FE, Lynch JP, Anderson CT, Ali JG, Felton GW (2022) Stomata-mediated interactions between plants, herbivores, and the environment. Trends in Plant Science 27: 287–300

Lu P, Liu J, Yu H, Li J (2026) MIRO1-mediated mitochondrial fusion is required for stomatal immunity in Arabidopsis. Nat Plants 12: 635–652

Ma J, Liang Z, Zhao J, Wang P, Ma W, Mai KK, Fernandez Andrade JA, Zeng Y, Grujic N, Jiang L, et al (2021) Friendly mediates membrane depolarization-induced mitophagy in Arabidopsis. Current Biology 31: 1931–1944.e4

Melotto M, Fochs B, Jaramillo Z, Rodrigues O (2024) Fighting for Survival at the Stomatal Gate. Annual Review of Plant Biology 75: 551–577

Melotto M, Underwood W, Koczan J, Nomura K, He SY (2006) Plant Stomata Function in Innate Immunity against Bacterial Invasion. Cell 126: 969–980

Módis K, Ju Y, Ahmad A, Untereiner AA, Altaany Z, Wu L, Szabo C, Wang R (2016) S-Sulfhydration of ATP synthase by hydrogen sulfide stimulates mitochondrial bioenergetics. Pharmacological Research 113: 116–124

Møller IM (2001) PLANT MITOCHONDRIA AND OXIDATIVE STRESS: Electron Transport, NADPH Turnover, and Metabolism of Reactive Oxygen Species. Annu Rev Plant Physiol Plant Mol Biol 52: 561–591

Murphy B, Bhattacharya R, Mukherjee P (2019) Hydrogen sulfide signaling in mitochondria and disease. The FASEB Journal 33: 13098–13125

Nietzel T, Elsässer M, Ruberti C, Steinbeck J, Ugalde JM, Fuchs P, Wagner S, Ostermann L, Moseler A, Lemke P, et al (2019) The fluorescent protein sensor ro GFP 2-Orp1 monitors *in vivo* H _2_ O _2_ and thiol redox integration and elucidates intracellular H _2_ O _2_ dynamics during elicitor-induced oxidative burst in Arabidopsis. New Phytol 221: 1649–1664

Pantaleno R, Schiel P, García-Mata C, Scuffi D (2024) Analysis of Guard Cell Readouts Using Arabidopsis thaliana Isolated Epidermal Peels. BIO-PROTOCOL. doi: 10.21769/BioProtoc.5033

Pantaleno R, Scuffi D, Costa A, Welchen E, Torregrossa R, Whiteman M, García-Mata C (2023) Mitochondrial H2S donor AP39 induces stomatal closure by modulating guard cell mitochondrial activity. Plant Physiology 191: 2001–2011

Pantaleno R, Scuffi D, García-Mata C (2021) Hydrogen sulphide as a guard cell network regulator. New Phytologist. 230: 451–456

Pantaleno R, Scuffi D, Schiel P, Schwarzländer M, Costa A, Garcia-Mata C (2025) Mitochondrial ß-Cyanoalanine Synthase Participates in flg22-Induced Stomatal Immunity. Plant Cell Environ. 48: 537–552. 10.1111/pce.15155

Papanatsiou M, Scuffi D, Blatt MR, García-Mata C (2015) Hydrogen Sulfide Regulates Inward-Rectifying K+ Channels in Conjunction with Stomatal Closure. Plant Physiology 168: 29–35

Pei D, Hua D, Deng J, Wang Z, Song C, Wang Y, Wang Y, Qi J, Kollist H, Yang S, et al (2022) Phosphorylation of the plasma membrane H+-ATPase AHA2 by BAK1 is required for ABA-induced stomatal closure in Arabidopsis. Plant Cell 34: 2708–2729

Qi J, Song C-P, Wang B, Zhou J, Kangasjärvi J, Zhu J-K, Gong Z (2018) Reactive oxygen species signaling and stomatal movement in plant responses to drought stress and pathogen attack: ROS signaling and stomatal movement. J Integr Plant Biol 60: 805–826

Qi J, Wang J, Gong Z, Zhou J-M (2017) Apoplastic ROS signaling in plant immunity. Current Opinion in Plant Biology 38: 92–100

Racca S, Gras DE, Canal MV, Ferrero LV, Rojas BE, Figueroa CM, Ariel FD, Welchen E, Gonzalez DH (2022) Cytochrome *c* and the transcription factor ABI4 establish a molecular link between mitochondria and ABA-dependent seed germination. New Phytologist 235: 1780– 1795

Racca S, Welchen E, Gras DE, Tarkowská D, Turečková V, Maurino VG, Gonzalez DH (2018) Interplay between cytochrome *c* and gibberellins during Arabidopsis vegetative development. Plant J 94: 105–121

Roussin-Léveillée C, Lajeunesse G, St-Amand M, Veerapen VP, Silva-Martins G, Nomura K, Brassard S, Bolaji A, He SY, Moffett P (2022) Evolutionarily conserved bacterial effectors hijack abscisic acid signaling to induce an aqueous environment in the apoplast. Cell Host & Microbe 30: 489–501.e4

Scuffi D, Álvarez C, Laspina N, Gotor C, Lamattina L, García-Mata C (2014) Hydrogen Sulfide Generated by L--Cysteine Desulfhydrase Acts Upstream of Nitric Oxide to Modulate Abscisic Acid-Dependent Stomatal Closure. Plant Physiology 166: 2065–2076

Scuffi D, Nietzel T, Di Fino LM, Meyer AJ, Lamattina L, Schwarzländer M, Laxalt AM, García-Mata C (2018) Hydrogen Sulfide Increases Production of NADPH Oxidase-Dependent Hydrogen Peroxide and Phospholipase D-Derived Phosphatidic Acid in Guard Cell Signaling. Plant Physiol 176: 2532–2542

Scuffi D, Pantaleno R, Schiel P, Niemeier J-O, Costa A, Schwarzländer M, Laxalt AM, García-Mata C (2025) Hydrogen Sulfide modulates Flagellin-Induced Stomatal Immunity. Plant Physiol 199: kiaf561

Shen J, Zhang J, Zhou M, Zhou H, Cui B, Gotor C, Romero LC, Fu L, Yang J, Foyer CH, et al (2020) Persulfidation-based Modification of Cysteine Desulfhydrase and the NADPH Oxidase RBOHD Controls Guard Cell Abscisic Acid Signaling. Plant Cell 32: 1000–1017

Szabo C, Ransy C, Módis K, Andriamihaja M, Murghes B, Coletta C, Olah G, Yanagi K, Bouillaud F (2014) Regulation of mitochondrial bioenergetic function by hydrogen sulfide. Part I. Biochemical and physiological mechanisms: Biochemistry of H _2_ S and mitochondrial function. Br J Pharmacol 171: 2099–2122

Telara Y, Akter S, Aroca A, Piccigallo L, Zhang D, Novi D, Lavilla-Puerta M, Gunawardana DM, La Monaca N, Lichtenauer S, et al (2025) Hydrogen sulfide modulates plant hypoxic responses through the persulfidation of Plant Cysteine Oxidases. doi: 10.1101/2025.11.05.686772

Vitvitsky V, Miljkovic JLj, Bostelaar T, Adhikari B, Yadav PK, Steiger AK, Torregrossa R, Pluth MD, Whiteman M, Banerjee R, et al (2018) Cytochrome *c* Reduction by H _2_ S Potentiates Sulfide Signaling. ACS Chem Biol 13: 2300–2307

Wagner S, Meyer AJ (2025) Illuminating plant metabolism with genetically encoded biosensors. Journal of Plant Physiology 311: 154498

Wang J, Xu G, Ning Y, Wang X, Wang G-L (2022) Mitochondrial functions in plant immunity. Trends in Plant Science 27: 1063–1076

Welchen E, Gonzalez DH (2016) Cytochrome *c*, a hub linking energy, redox, stress and signaling pathways in mitochondria and other cell compartments. Physiol Plantarum 157: 310–321

Welchen E, Hildebrandt TM, Lewejohann D, Gonzalez DH, Braun H-P (2012) Lack of cytochrome c in Arabidopsis decreases stability of Complex IV and modifies redox metabolism without affecting Complexes I and III. Biochimica et Biophysica Acta (BBA)-Bioenergetics 1817: 990–1001

Yamaguchi Y, Nakamura T, Kusano T, Sano H (2000) Three Arabidopsis Genes Encoding Proteins with Differential Activities for Cysteine Synthase and-Cyanoalanine Synthase. Plant and Cell Physiology 41: 465–476

Yang Y, Zhao Y, Zhang Y, Niu L, Li W, Lu W, Li J, Schäfer P, Meng Y, Shan W (2022) A mitochondrial RNA processing protein mediates plant immunity to a broad spectrum of pathogens by modulating the mitochondrial oxidative burst. The Plant Cell 34: 2343–2363

Yang Y, Zhao Y, Zhao W, Zhang Y, Wang H, Grant M, Schäfer P, Meng Y, Shan W (2025) Mitochondrial ROS trigger interorganellular signaling and prime ER processes to establish enhanced plant immunity. Sci Adv 11: eady9234

Yuan M, Ngou BPM, Ding P, Xin X-F (2021) PTI-ETI crosstalk: an integrative view of plant immunity. Current Opinion in Plant Biology 62: 102030

Zhou H, Zhou Y, Zhang F, Guan W, Su Y, Yuan X, Xie Y (2021) Persulfidation of Nitrate Reductase 2 Is Involved in l-Cysteine Desulfhydrase-Regulated Rice Drought Tolerance. IJMS 22: 12119

Zipfel C, Robatzek S, Navarro L, Oakeley EJ, Jones JDG, Felix G, Boller T (2004) Bacterial disease resistance in Arabidopsis through flagellin perception. Nature 428: 764–767

