## Supplementary Material Pantaleno for "Mitochondrial H2S and cytochrome c coordinate guard cell redox dynamics during stomatal immunity"

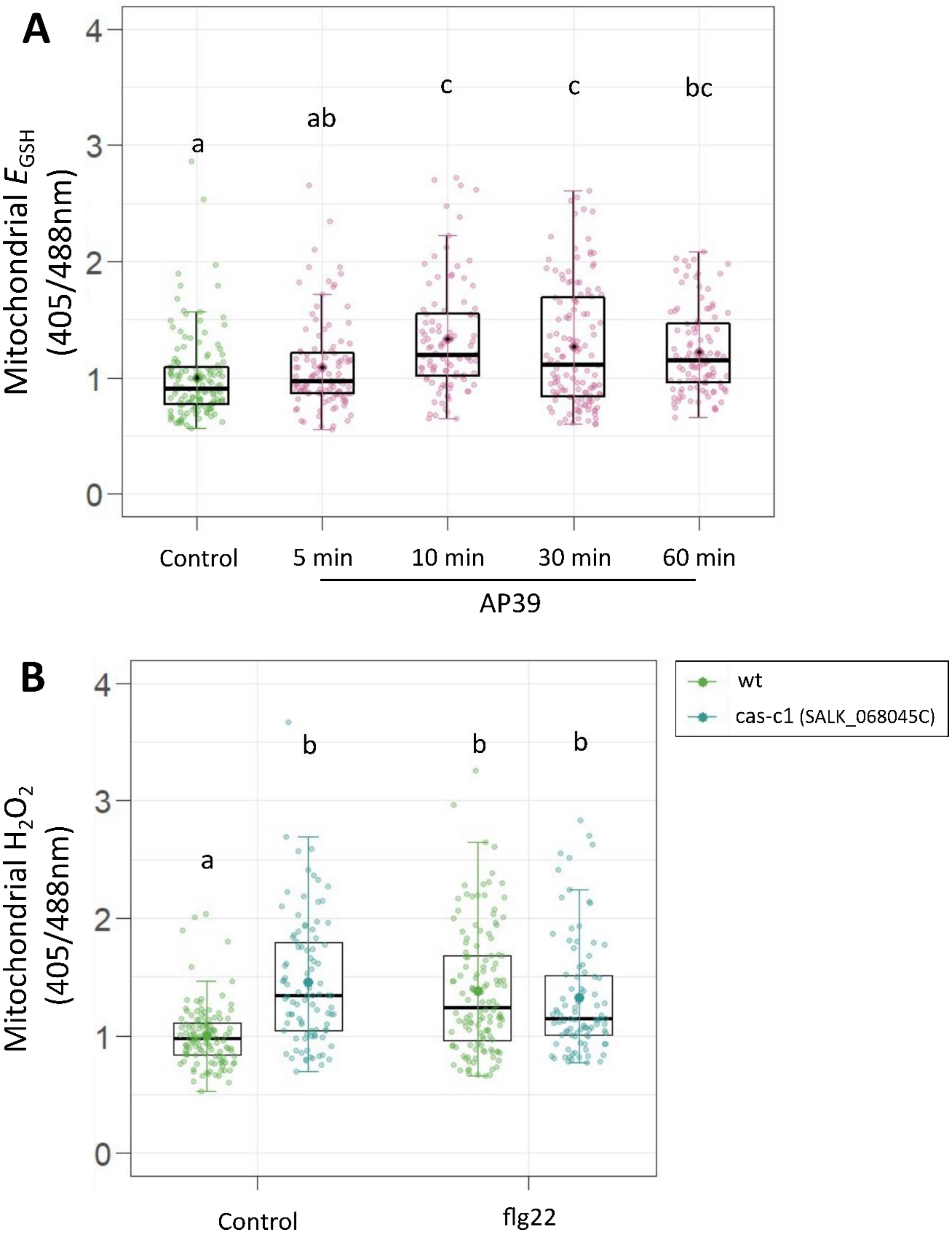


**Supplementary Figure S1: AP39 and flg22 modulates mitochondrial redox state in guard cells.** Epidermal strips from *Arabidopsis thaliana* wild-type (wt) leaves expressing the mt-Grx1-roGFP2 biosensor for mitochondrial $E_{GSH}$ detection (A) or wt and *cas-c1* (SALK_ 068045C) expressing mt-roGFP2-Orp1 to monitor mitochondrial $H_2O_2$ (B), were incubated in opening buffer (5 mM MES,

50 mM KCl, pH = 6.1) for 7 hr and then treated with: 100 nM AP39 for 5, 10, 30 and 60 min (A) or 1 µM flg22 for 5 min (B). Guard cells fluorescence was detected by confocal microscopy, and 405/488 nm ratios were calculated for each guard cell, normalized to the control of wt and represented as individual points. Boxplots are bound by the 25th to 75th percentile. The line in the middle is the median, the darker point is the mean and whiskers span 10th to 90th percentile. Black points represent the outliers. Data is from at least three independent replicates for each experiment. Different letters indicate statistical differences among treatments (Tukey's method, confidence level = 0.95).

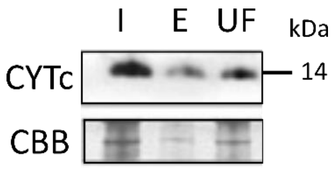


**Supplementary Figure S2: CYTc is persulfidated *in vivo*.** Western blot analysis of persulfidated proteins purified by dimedone switch method (I: input, E: eluate, and UF: unbound fraction). An anti-CYTc (CYTc) antibody was used to detect CYTc, and Coomassie Brilliant Blue (CBB) staining was used as a loading control.

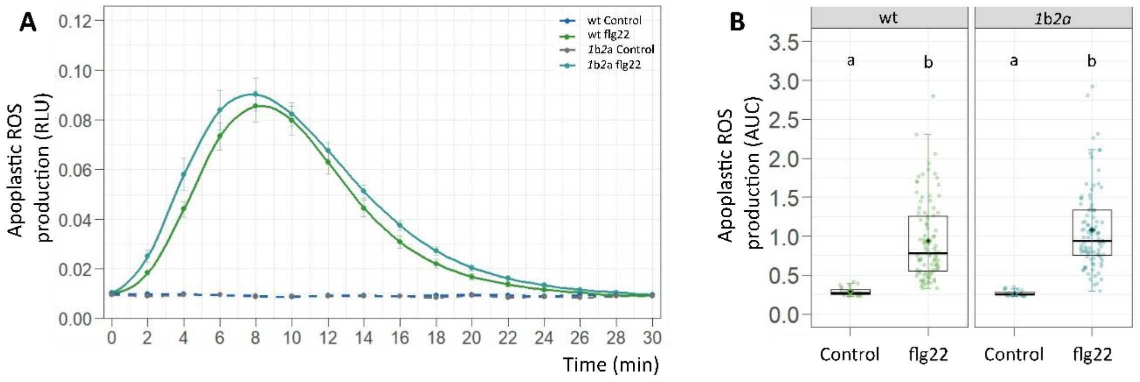


**Supplementary Figure S3: Apoplastic ROS production in *cytc* mutants.** Leaf discs from 5-week-old *Arabidopsis thaliana* wild-type (wt) or *cytc* double mutant (*1*b*2*a) plants were floated in deionized water overnight. The following day, they were placed on 96-well white plate containing a solution with 20 mM luminol and 0.02 mg/mL horseradish peroxidase. ROS production was triggered with 100 nM flg22 and luminescence was measured with a luminometer. Apoplastic ROS production is expressed as the relative light units (RLU) detected (A), or the area under the curve (AUC) for each leaf disc (B). In (A) the points represent the mean RLU over time and the error bars are the standard error. In (B) the AUC are represented as points in the boxplots, where the box is bound by the 25th to 75th percentile. The line in the middle is the median, the darker point is the mean and whiskers span 10th to 90th percentile. Black points represent measurements outside those margins. Data was taken from at least three independent experiments. Different letters indicate statistical differences among treatments (Tukey's method, confidence level = 0.95).
